# Whole-exome analysis in Parkinson’s disease reveals a high burden of ultra rare variants in early onset cases

**DOI:** 10.1101/2020.06.06.137299

**Authors:** Bernabe I. Bustos, Dimitri Krainc, Steven J. Lubbe, for The International Parkinson’s Disease Genomics Consortium (IPDGC)

## Abstract

Parkinson’s disease (PD) is a complex neurodegenerative disorder with a strong genetic component. We performed a “hypothesis-free” exome-wide burden-based analysis of different variant frequencies, predicted functional impact and age of onset classes, in order to expand the understanding of rare variants in PD. Analyzing whole-exome data from a total of 1,425 PD cases and 596 controls, we found a significantly increased burden of ultra-rare (URV= private variants absent from gnomAD) protein altering variants (PAV) in early-onset PD cases (EOPD, <40 years old; P=3.95×10^−26^, beta=0.16, SE=0.02), compared to LOPD cases (>60 years old, late-onset), where more common PAVs (allele frequencies <0.001) showed the highest significance and effect (P=0.026, beta=0.15, SE=0.07). Gene-set burden analysis of URVs in EOPD highlighted significant disease- and tissue-relevant genes, pathways and protein-protein interaction networks that were different to that observed in non-EOPD cases. Heritability estimates revealed that URVs account for 15.9% of the genetic component in EOPD individuals. Our results suggest that URVs play a significant role in EOPD and that distinct etiological bases may exist for EOPD and sporadic PD. By providing new insights into the genetic architecture of PD, our study may inform approaches aimed at novel gene discovery and provide new directions for genetic risk assessment based on disease age of onset.

## INTRODUCTION

Parkinson’s disease (PD) is a complex neurodegenerative disorder characterized by the loss of dopaminergic neurons in the substantia nigra (SN), leading to motor dysfunction and a progressive neurodegenerative syndrome. Several common and rare disease-associated genes and loci have been discovered in families and large case-control studies; however, they currently explain only about 26-36% of the disease heritability^1^, therefore additional novel variants and genes associated with PD are yet to be found.

Despite the recent explosion of available genetic data for PD, the vast majority of cases still have an unknown genetic basis^2^. The use of next generation sequencing (NGS) in PD has enabled large-scale whole-exome sequencing (WES) studies in both families and case-controls unrelated populations^3^. Several studies have leveraged the use of WES to analyze candidate genes or known biological pathways and their load of rare variants^4-8^. However, to date no study has systematically evaluated the burden of rare (minor allele frequency <0.01) damaging variants in a “hypothesis-free” approach with functional insights, that could help to expand the genetic architecture of PD, early-onset PD (EOPD) and late-onset PD (LOPD), in order to potentially uncover novel susceptibility genes and pathways.

In the present study, we used two case-control WES cohorts from the International Parkinson’s disease Genomics Consortium (IPDGC, www.pdgenetics.org)^5^ and the Parkinson’s Progression Markers Initiative (PPMI, www.ppmi-info.org)^9^, and conducted a systematic assessment of the burden of variants across different allele frequencies, functional impact predictions and different ages of onset (AOO) at the whole-exome, gene-set and gene-wise levels in idiopathic PD cases compared to healthy controls. Using a large compendium of functional gene-sets, molecular pathways, tissue and cell-type specific gene expression profiles, and protein-protein interaction (PPI) networks, our results revealed a significant burden of damaging ultra-rare variants (URVs) in EOPD cases that are enriched in disease-relevant molecular pathways, and suggest major differences in the genetic and functional architecture compared to LOPD.

## RESULTS

### Exome-wide contribution of rare damaging coding variants to Parkinson’s disease

We systematically assessed the contribution of coding variants to PD in two independent case-control cohorts of European descent: (i) IPDGC^5^, comprising 1,042 PD cases and 452 controls, and (ii) PPMI^9^, with 383 PD cases and 144 controls (See complete pipeline in **Supplementary Fig. 1**). We also stratified cases according to AOO (**Table 1**). The IPDGC cases had a mean AOO of 43.02 years (SD=10.63), including 438 EOPD (AOO <40 years of age) and 604 PD cases above 40 years of age (40+). Controls had average age of recruitment of 46.20 years (SD=26.07). The PPMI cases had a mean AOO of 61.62 years (SD=9.72), and comprised 9 EOPD and 374 40+ (including 232 LOPD individuals). Controls had an average age of recruitment of 61.11 years (SD=10.18). Average per sample transitions/transversions (Ts/Tv), homozygous/heterozygous (Hom/Het) and insertions/deletions (In/Del) ratios were among expected ranges for both cohorts, respectively (IPDGC=3.2, 1.69, 0.87; PPMI=3.19, 1.82, 0.69).

**Table 1.** Whole-exome cohorts analyzed in the present study.

| Whole-exome cohort | Median age of PD onset | Median age of recruitment in controls | Burden groups | Sample size | Cases | Controls |
| --- | --- | --- | --- | --- | --- | --- |
| IPDGC | 43.02 (± 10.63) | 46.20 (± 26.07) | All | 1,494 | 1,042 | 452 |
|  |  |  | Cases <40 y/o (EOPD) | 890 | 438 | 452 |
|  |  |  | Cases >40 y/o | 1,056 | 604 | 452 |
| PPMI | 61.62 (± 9.72) | 61.11 (± 10.18) | All | 527 | 383 | 144 |
|  |  |  | Cases >40 y/o | 518 | 374 | 144 |
|  |  |  | Cases >60 y/o (LOPD) | 376 | 232 | 144 |

We classified and counted alleles according to frequency, functional and deleteriousness predictions, in different AAOs in cases and controls (average counts on **Supplementary Table 1**). With regression analyses we observed a highly significant burden of all three classes of ultra-rare variants in the IPDGC EOPD cases compared to controls (URVs= singleton variants in our cohort, having an allele count of 1/2988 and without reported frequency in gnomAD^10^): Protein altering variants (PAV), P=3.95×10^−26^, beta=0.158, SE=0.015; Nonsynonymous (NSN), P=7.36×10^−26^, beta=0.162, SE=0.015; and Loss-of-Function (LoF), P=6.1×10^−4^, beta=0.295, SE=0.086 **(Fig. 1A, Supplementary Table 2)**. LoF variants appeared to have a larger contribution compared to NSN variants in EOPD cases. Similar but weaker results were observed for all classes of URVs in all PD cases, except for LoF variants that did not surpass Bonferroni correction (PAV, P=9.17×10^−18^, beta=0.102, SE=0.012; NSN, P=2.09×10^−17^, beta=0.105, SE= 0.012; LOF, P=0.004, beta=0.209, SE=0.073) as well as in 40+ PD cases (PAV, P=1.92×10^−6^, beta=0.065, SE=0.014; NSN, P=2.75×10^−6^, beta=0.066, SE=0.014; LOF, P=0.098, beta=0.133, SE=0.08). After removing lower frequency variants for each category, we observed that the top burden results were for more common allele frequencies (*i.e*. > singletons) in 40+ PD cases (P=0.003, beta=0.01, SE=0.003; **Fig. 1B, Supplementary Table 3**), although this did not survive multiple testing correction. We next analyzed synonymous and noncoding variants. While we did not observe increased enrichment across the majority of frequencies and effect classes (“Damaging”, CADD >12.37; “Benign”, CADD <12.37), we observed significant URV association signals surviving multiple testing correction in EOPD for damaging and benign synonymous variants (P=9.13×10^−16^, beta=0.38, SE=0.048; P=1.80×10^−15^, beta=0.22, SE=0.028, respectively), and benign noncoding variants (P=6.37×10^−8^, beta=0.21, SE=0.038; **Supplementary Fig. 2, Supplementary Table 4**). Similar associations were observed for all PD cases and only damaging synonymous variants for 40+ PD. We next assessed if our URV burden results were biased by an artificial excess of rare variants due to sequencing errors or other hidden confounding factors. We therefore corrected the burden tests for URVs by the total number of singleton variants (regardless of their frequency in gnomAD), and observed that the associations remained significant, we even noted increases in the significance and effect size estimates after correction (URVs not corrected, P=2.27×10^−19^, beta=0.06, SE=0.006; corrected, P=1.43×10^−26^, beta=0.08, SE=0.008). Using a stricter read depth threshold (DP>20), we reevaluated URVs burden across frequencies and functional annotations, and observed that association signals remained highly significant (**Supplementary Table 5**). To further explore increased burden of coding variants in PD, we next analyzed the PPMI cohort. Although no results survived Bonferroni correction, we observed the highest burden for PAVs with frequency <0.001 in all PD cases (P=0.01, beta=0.02, SE=0.009). Interestingly, the strongest effect size was seen for LoFs with frequency <0.001 in LOPD (P=0.026, beta=0.152, SE=0.07) (**Supplementary Fig. 3A, Supplementary Table 6)**. To explore this further, we repeated the analysis after removing URVs and observed that both the effect size and significance increased (P=0.014, beta=0.20, SE=0.08; **Supplementary Fig. 3B, Supplementary Table 7**). For synonymous and noncoding variants, no frequency category survived multiple testing correction (**Supplementary Fig. 4, Supplementary Table 8**).

**Figure 1.**
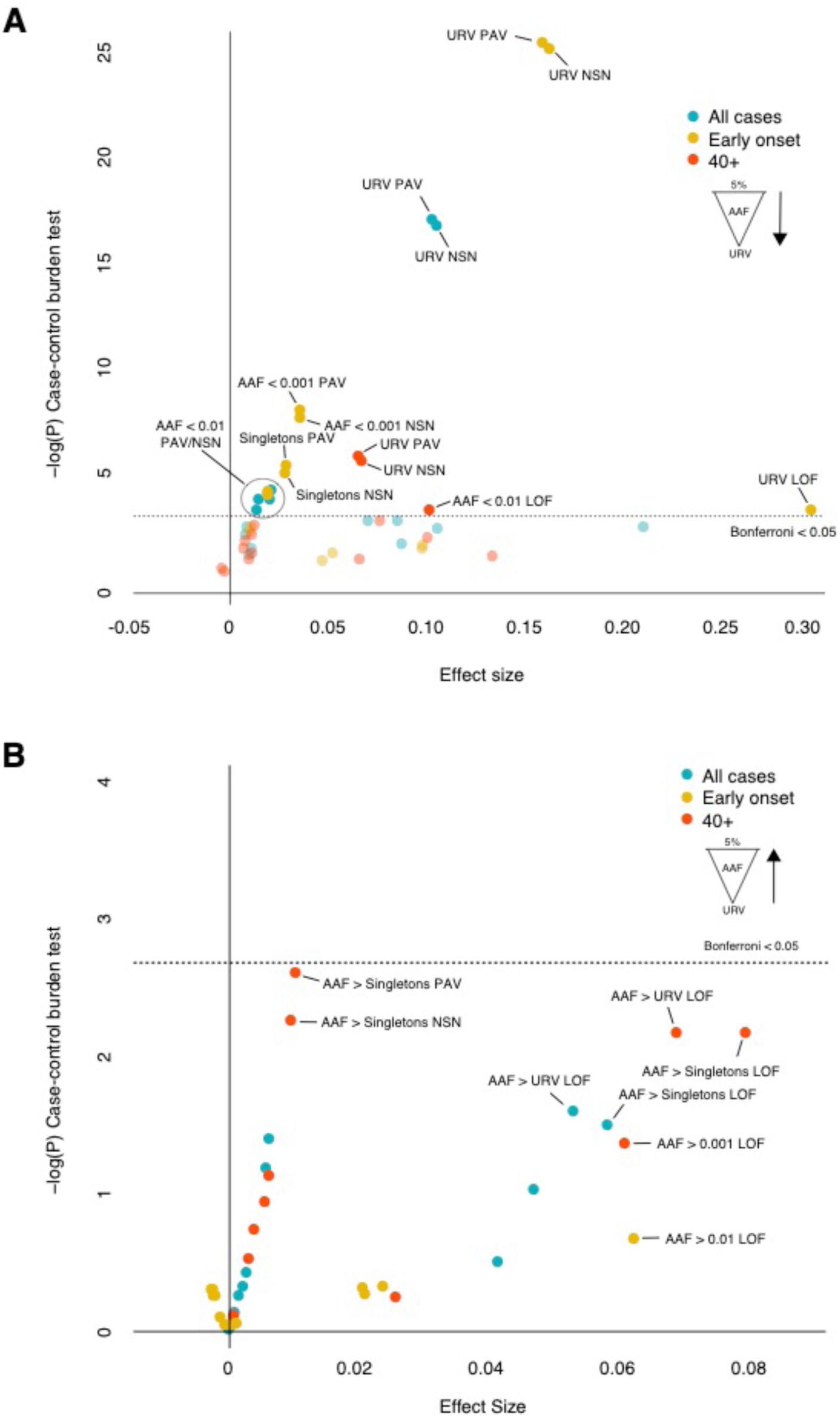
Whole-exome burden of rare coding damaging variants in the IPDGC cohort. Burden analysis of different variant frequencies and functional categories for all PD cases, cases over under 40 years old (EOPD) and over 40 years old (40+). Logistic regression -log_10_(P-values) and beta coefficients are represented for: **A)** Burden from variants with 5% or less frequency to ultra-rare variants (URVs). **B)** Burden towards more common variants removing lower frequency variants from each variant category. AAF: Alternative allele frequency; PAV: Protein-altering variants; NSN: Nonsynonymous variants; LoF: Loss-of-function variants.

### Functional landscape of PD-associated damaging coding variants

#### Enrichment in gene-sets of highly constrained genes

Using the Sequence Kernel Association Test–Optimal (SKAT-O)^11^, we observed the most significant enrichment for NSN URVs within genes with high probability of being intolerant to heterozygous loss-of-function mutations (pLI >0.9, P=1.25×10^−20^) followed by genes highly intolerant to 2 loss-of-function mutations (recessive) (pRec >0.9, P=1.74×10^−18^) and lastly in genes highly tolerant to loss-of-function variation (pNull >0.9, P=1.35 ×10^−9^) **(Fig. 2A)**. For PAVs, the most significant enrichment was observed in pRec >0.9 (P=2.22×10^−17^), followed by pLI >0.9 (6.42×10^−17^) and then pNull >0.9 (P=1.65×10^−11^). LoF variants showed enrichment only in pRec >0.9 (P=0.01) and pNull >0.9 gene-sets (P=0.02). For IPDGC 40+ PD cases, we analyzed the variants with frequency above singletons and observed a significant enrichment of PAVs and NSNs variants across all three categories, with the most significant signal in pRec >0.9 genes (PAV, P=1.55×10^−5^; NSN, P=3.0×10^−4^; **Supplementary Fig. 5A**). For the PPMI cohort, no significant associations were observed after multiple testing correction (**Supplementary Table 9**); however, the top signals P-values were observed in LoFs with frequency above rare singleton in pRec >0.9 genes analyzed in 40+ PD cases, and in LoFs with frequency <0.001 without URVs in pNull >0.9 genes analyzed in LOPD cases.

**Figure 2.**
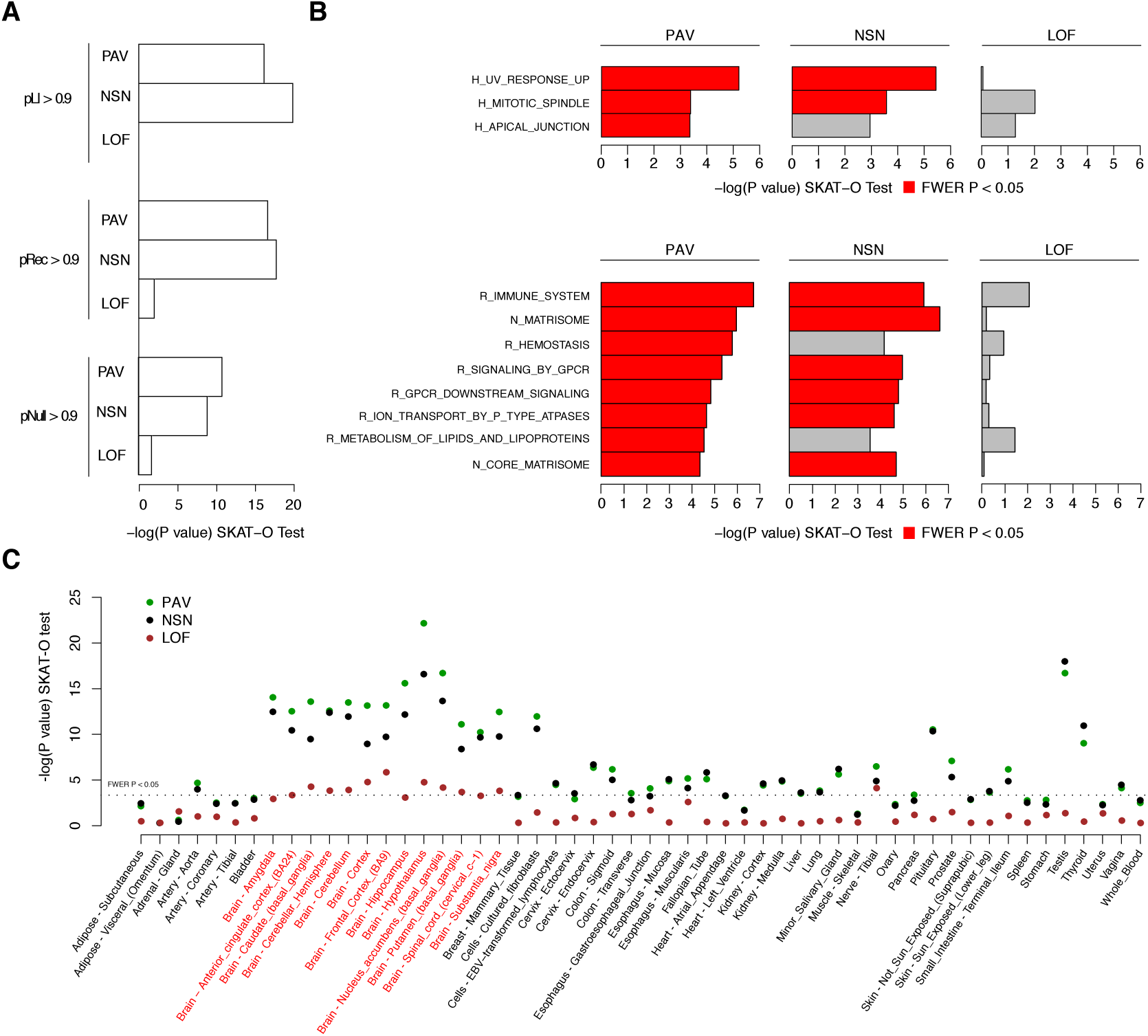
Gene-set burden analysis in the IPDGC cohort for EOPD cases. Gene-set analysis performed with SKAT-O using PAV, NSN and LoF URVs using gene lists from: **A)** gnomAD gene constraint metrics pLI, pRec and pNull (>0.9); **B)** MsigDB Hallmark pathways (upper panel) and C2 curated pathways (lower panel); **C)** High expression genes from 54 tissues in the GTEx database V8. A family-wise error rate P <0.05 was used as a threshold for statistically significant results.

#### Enrichment in molecular gene-sets and pathways

Using the Hallmark MsigDB^12^ gene-sets, we observed significant enrichment in NSN URVs within genes up-regulated in response to ultra-violet radiation (UV) (MsigDB name=M5941, P=3.58×10^−6^; **Fig. 2B upper panel**), followed by the Mitotic Spindle (M5893, P=2.68×10^−4^). For PAVs, we observed enrichment in genes found in the Apical Junction pathways (M5915, P=4.46×10^−4^). In the C2 gene-sets, the top enrichment was seen for PAV URVs within the Immune System pathway (M1045, P=1.84×10^−7^, **Fig. 2B lower panel**) and the Reactome Metabolism of Lipids and Lipoproteins (M27451, P=2.86×10^−5^). For NSN URVs, the NABA Matrisome pathway (M5889, P=2.28×10^−7^) was observed to be enriched. For less rarer variants in IPDGC 40+ PD cases (frequency > singletons), the top enriched Hallmark pathway was for NSNs within the Bile Acid Metabolism (M5948, P=1.63×10^−8^), and for the C2 gene-set, the NABA extra cellular membrane Affiliated Proteins pathway (M5880, P=6.03×10^−8^). Different pathways were highlighted between the Hallmark and C2 gene-sets, with the only overlap found in the C2 NABA Matrisome pathway (M5889, P=4.67×10^−6^, **Supplementary Fig. 5B lower panel**). No significant results were observed for variants in 40+ PD (frequency > singletons) and LOPD (frequency <0.001 without URVs) in the PPMI dataset (**Supplementary Table 10**). Previous reports that had performed pathway burden analyses have found enrichment of rare damaging NSN variants in both lysosomal storage disorder^7^ (LSD) genes (frequency <0.03) and mitochondrial DNA maintenance pathway genes (frequency <0.01)^13^. To validate our hypothesis-free methodology, we performed burden analysis on these two specific pathways. We found that URVs were significantly enriched in the mitochondrial DNA maintenance on EOPD individuals (P=0.016). We also observed that the LSD genes enrichment was only significant when using more common variants (frequency <0.05) and that the association was driven by 40+ PD cases (P=1.67×10^−4^, **Supplementary Table 11**).

#### Enrichment in highly expressed genes across specific tissues from GTEx

We observed significant enrichment signals for genes harboring URVs in EOPD individuals in all GTEx brain tissues^14^, for all three types of variant categories (**Fig. 2 C**). Interestingly, significant enrichment of PAVs and NSN variants was also detected in several other tissues. For 40+ PD cases, we observed a different landscape for PAVs and NSNs, with no enrichment surpassing multiple testing correction in any brain tissue. For LoF variants the enrichment was significant in other tissues suggesting a distinct contribution compared to the observations in EOPD (**Supplementary Fig. 5C**). No significant results were observed in the PPMI dataset (**Supplementary Table 12**).

### Protein-protein interaction networks and single-cell enrichment analysis of gene-wise burden

#### Gene-wise burden and PPI analysis

Although no genes survived multiple-testing correction, we carried out a protein-protein interaction (PPI) network analysis to explore all nominally significant genes (unadjusted SKAT-O P<0.05, n=308; **Supplementary Table 13**) in EOPD and 40+ PD cases. Leveraging both WebgestaltR^15^ and STRING^16^, we observed a significant network (STRING P=0.01, **Supplementary Fig. 6**). No significant gene ontology (GO) enrichment was observed, however, the top GO term detected was the Regulation of Microtubule Cytoskeleton Organization (GO:0070507) which was driven by nine URV containing genes: *AKAP9, APC, CAMSAP2, CDK5RAP2, CEP120, CKAP2, CYLD, KIF11* and *TPR* **(Supplementary Table 14)**. After including 25 known and suggested PD genes^17^ (**Supplementary Table 15**), the network significance dramatically increased (STRING P<1×10^−16^, **Fig. 3A**). Within this network, we observed significant GO enrichment within the Negative Regulation of Neuron death pathway (GO:1901215, FDR=4.93×10^−6^; **Supplementary Table 16**), containing 7 genes harboring URVs: *FGF8, HYOU1, ITSN1, PPP5C, MAP3K5, NAE1*, and *SIRT1*. We next repeated this analysis, but now including 303 genes tagged by the 90 PD GWAS loci^1^, which also yielded a significant network (STRING P <1×10^−16^, **Supplementary Fig. 7**). Within this network, the most significant specific GO biological process was the Intrinsic apoptotic signaling pathway in response to endoplasmic reticulum (ER) stress (GO:0070059, FDR=4.2×10^−3^, **Supplementary Table 17**) containing five genes harboring URVs (*CREB3, ERN1, HYOU1, MAP3K5*, and *SIRT1*).

**Figure 3.**
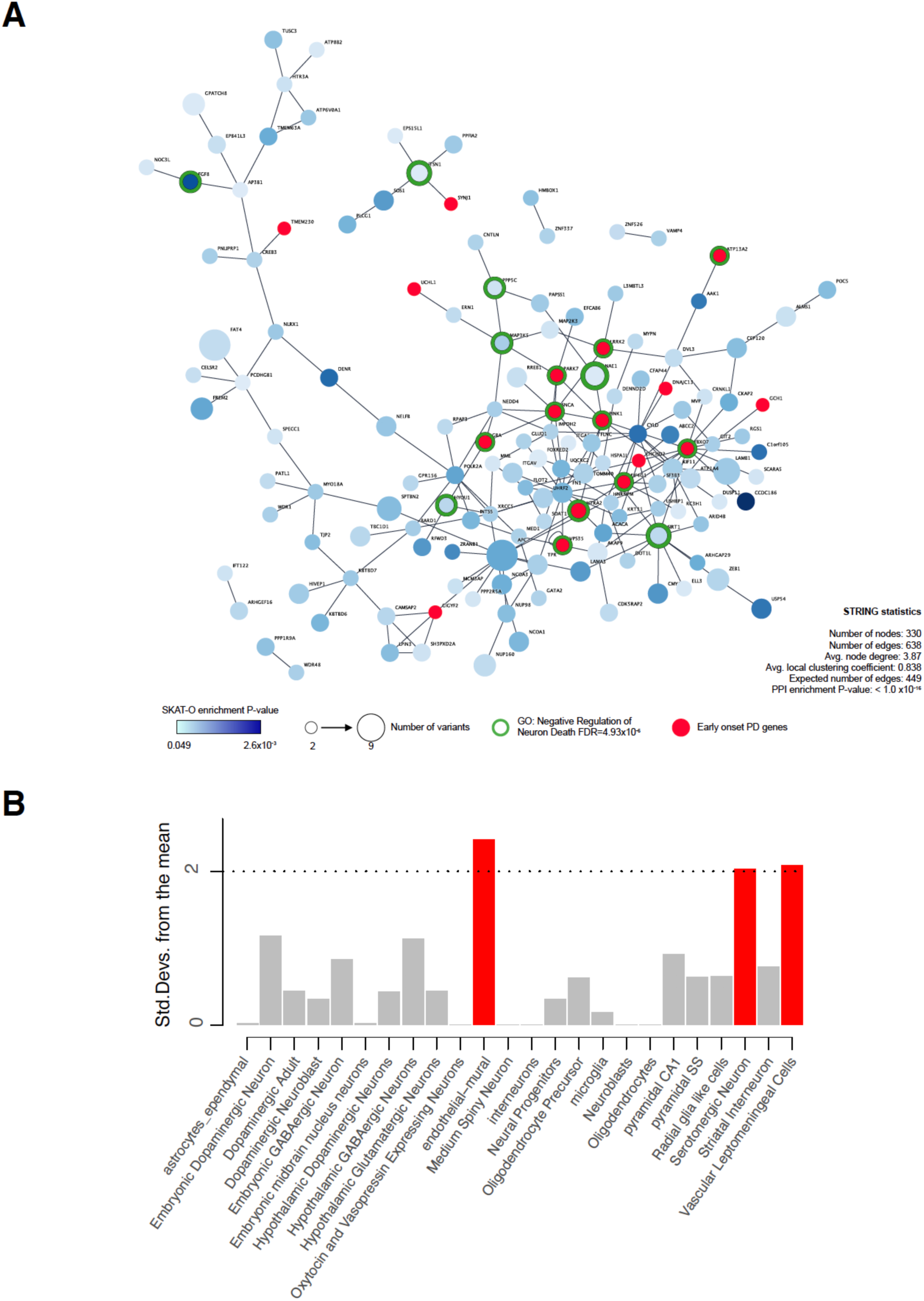
Protein-protein Interaction Network and Single-Cell enrichment analysis of gene-wise burden results in EOPD. **A)** Network built using 308 genes with SKAT-O P <0.05 from EOPD URVs and 26 known and suggested PD causal genes with WebGestalt Network Topology-based Analysis (NTA). Statistics from STRING using the same list of genes are included. Shades of blue indicate SKATO P-values from less (light-blue) to more significant (dark-blue) burden association. Node size indicate the number of variants observed in each gene and the green perimeter around the nodes show genes members of significant gene-ontology enrichment results. **B)** Single-cell and single-nuclei RNASeq enrichment analysis of EOPD gene-wise burden list leveraged using expression datasets from 24 mice brain tissues. Red bars represent significant results surpassing a FDR cutoff of <0.05.

#### Enrichment in specific brain cell types

We next aimed to identify specific mouse brain cell-types expressing the 308 URV harboring genes using EWCE^18,19^. We observed significant enrichment on the following three cell types (FDR P <0.05): serotonergic neurons; endothelial mural cells, and vascular leptomeningeal cells (**Fig. 3 B**). These results show that URVs could be playing an important role in specific cell-types within the brain.

### Heritability of URVs in PD

Heritability estimation in EOPD using SingHer^20^, showed that URVs account for 15.9% of the genetic component, 4.6% to All PD cases and 2.2% to 40+ PD. These results support our previous findings regarding the difference of URVs between different AOOs in PD.

## DISCUSSION

Here we have comprehensively studied the burden of rare variants in PD using WES datasets from 1,425 idiopathic PD cases and 596 healthy controls. We have found an increased burden of damaging URVs across affected cases, and observed the highest significance in EOPD individuals, suggesting that some of the missing heritability of EOPD may be accounted for by private variants. URVs have been increasingly gaining attention due to the explosion of NGS-based studies, and have been shown to have a major impact in gene-expression regulation, explaining near ∼25% of the heritability^20^. Similar increased URVs burdens have been seen in schizophrenia^21^, epilepsy^22^ and Alzheimer’s disease^23^, being identified as an important source of genetic risk. Here, we estimated that URVs contribute ∼15% towards the heritability of PD which is more than half of the current heritability estimates derived from GWAS identified common variants (∼26%)^1^. This indicates that URVs may play a major sizeable role in PD etiology thereby warranting further research.

Unique patterns of constrained gene-set enrichment were observed for EOPD and 40+ PD/LOPD. URVs in EOPD predominantly mapped to highly constrained genes (pLI >0.9) suggesting that EOPD may be driven by hetero- and homozygous highly deleterious private variants/URVs. In contrast, 40+ PD and LOPD appear to be driven by more common/infrequent variants (frequency >rare singletons) in genes that are intolerant to homozygous LoF variants mostly (pRec >0.9). Their presence within the general population suggests that these variants may be less damaging or have incomplete penetrance, further complicating variant discovery efforts. Our results therefore support previous assertions that different genes/variants and inheritance patterns contribute to EOPD and LOPD^24^. While EOPD-specific molecular signatures have recently been uncovered^25^, more studies are needed to investigate the different etiologies in EOPD and LOPD.

While it is premature to comment on the functional pathways highlighted in the 40+ PD and LOPD, many of the pathways pulled out in the EOPD analysis are of interest. Much research has investigated the proposed genetic link between PD, pigmentation (peripheral melanin and neuromelanin) and melanoma^26,27^, therefore the identified high burden of EOPD URVs in the MsigDB hallmark UV response pathway further highlights the link between PD and melanoma, and additionally implicates a role for pigmentation in disease pathology. The mitotic spindle pathway is associated with microtubule stabilization, a process known to be dysregulated in PD^28^. Recent evidence has shown that dopamine deficiency (a hallmark of PD) perturbs circadian/mitotic gene networks resulting in increased expression of mitotic spindle pathway genes in the striatum of PD patients^29^. Our observations of increased enrichment of URVs in genes involved in this pathway adds support to its potential role in PD etiology. The apical junction pathway is composed of genes implicated in cell-cell contacts, which are structurally important for different biological functions, such as the intestinal epithelial barrier^30^. Structural alterations of the intestinal epithelial barrier have been found in PD patients^31^, and in Crohn’s disease – both of which have *LRRK2* as common genetic factor^32^. Thus, our findings of high burden of URVs in the apical junction pathway may provide some evidence for the observed link between the etiology of PD and the gastrointestinal track. URVs enriched in pathways relating to the Immune System and the Metabolism of Lipids and Lipoprotein are interesting targets for functional validation, since they are established biological processes affected in PD pathophysiology^33,34^. The enrichment observed in the Matrisome pathway is of note, since constituent genes are key components of the extra cellular matrix (ECM). Functional and structural damage to the ECM during aging has been observed^35^, impacting important brain structures such as the blood-brain barrier^36^, which is known to be affected in both PD and Alzheimer’s disease^37^. The distinct pattern of enrichment of URV harboring genes in brain tissues provide some evidence of their possible functional role, along with the pathways found. Intriguingly, there is again no overlap in the enrichment in EOPD compared to 40+ PD cases, showing different pathways which reinforce the notion of distinct genetic architectures between AOOs. It is of note that for 40+ PD cases, the enrichment found in non-brain regions suggests that the roles of the genes/variants involved in LOPD pathogenesis may not be restricted to the brain, which is in agreement with recent evidence indicating that PD risk loci genes do not lie only in specific brain cell types or regions, but are involved in global cellular processes detectable across other tissues/organs^38^.

Although not a single gene survived multiple testing correction, several genes with suggestive enrichment of URVs are interesting candidates. *SLC39A1* mutations have been found to disrupt manganese homeostasis and cause childhood-onset parkinsonism-dystonia^39^. *SRGAP3* is differentially expressed in dopaminergic SN neurons^40^, and it is part of the axon guidance pathway already implicated in PD^41,42^. *FGF8* improves dopaminergic cell survival and functional restoration in a rat PD model^43^. *HYOU1* plays an important role in hypoxia-induced apoptosis as it accumulates within the endoplasmic reticulum (ER) under hypoxic conditions^44^ with its suppression associated with accelerated cell death^45^. The significant PPI networks identified here (GO:1901215, 0070059), while adding strength to our approach, further highlight the important roles that neuronal death regulation and ER stress play in PD etiology. Much research has linked ER stress response to PD pathogenesis (reviewed in Colla, 2019^46^), and pathways related to ER stress such as the unfolded protein response are strongly differentially expressed in prefrontal cortex of PD patients^47^. Several highly interconnected genes within these networks are of interest: *SIRT1* (11 interactions) has been shown to be protect against PD in cellular and animal models^48^; *MAP3K5* (4 interactions) is inhibited by *DJ-1* (PARK7)^49^; and *ITSN1* (4 interactions) is involved in endosomal and lysosomal trafficking^50^, which contributes to PD risk through increased expression^51^.

At the cell-type level, the significant enrichment observed in endothelial-mural cells (vascular smooth muscle cells and pericytes that constitute key structures of the blood-brain barrier^52^) is in line with our observations of high burden of URVs in the Matrisome pathway. The observed enrichment in serotoninergic neurons gives novel insights into previous genetic observations of serotonergic dysfunction in PD, where *SNCA* p.Ala53Thr carriers have reduced brain serotonin transporters^53^ and *LRRK2* mutation carriers have increased serotonin transporters in the brain^54^. Enrichment in vascular leptomeningeal cells in conjunction with recent GWAS findings^55^, suggests that both common and URVs within genes important for brain repair mechanisms and neurogenesis^56^ play a role in PD pathology. Together, these data help us expand our understanding of the disease etiology and the role of URVs in PD pathogenesis.

The main limitation of this work relates to insufficient statistical power to detect variant burden and the lack of appropriate age-matched replication cohorts, despite the fact that the IPDGC and PPMI PD cohorts are some of the largest WES datasets available. Not only do we expect that larger genetic cohorts will help replicate our findings, they would also edge this study closer to the prohibitively large sample sizes that are currently needed to achieve the sufficient power. Our observations that both synonymous and noncoding variants appear enriched in PD suggest that our findings could be inflated by a global spurious excess of singleton variants within our datasets. However, our strict QC (Ts/Tv ratios within acceptable ranges) and the fact that after correcting the URV burden for the total number of singletons our effect sizes increased and gained significance argue against this. Synonymous and noncoding variants are known to contribute to disease pathology through a wide variety of mechanisms, mostly associated to the regulation of gene expression through disruption of transcription factor binding sites, enhancers and splicing sites^57-59^. It is therefore plausible that these variants are not completely neutral and may contribute to PD etiology, however, more studies are needed to accurately elucidate their role in PD. Overall, this study shows that URVs have a major contribution to PD, especially in early-onset individuals which have a distinct genetic background from sporadic cases. The identified URVs are significantly enriched in disease relevant gene-sets, pathways and genes that interact with each other as well as with known and suggested PD genes. Our results help to broad the understanding of PD genetic risk load in different age of onset and expand the landscape of biological pathways potentially involved in this devastating disease.

## METHODS

### Ethical Statement

This study was approved by the corresponding local ethical scientific committees. More detailed data usage authorizations are provided elsewhere^5,9^.

### Samples used in the study

#### International Parkinson’s Disease Genomics Consortium (IPDGC) WES cohort

The IPDGC WES data used consisted of a total number of 1,933 self-reported European individuals, composed of 1,398 PD cases and 535 neurologically healthy controls (https://pdgenetics.org/resources). A more detailed description of the cohort is reported elsewhere^5^.

#### Parkinson’s Progression Markers Initiative (PPMI) WES study

The PPMI WES dataset includes 645 individuals, composed of 462 PD cases and 183 healthy controls, all of self-reported European descent. Data were obtained from the PPMI database under the appropriate PI membership (www.ppmi-info.org/data)^9^.

### Data processing and quality controls

WES data generation and processing details for both IPDGC and PPMI cohorts have been previously described^5,9^. All analyses were performed using VCF files acquired through authorized requests.

For both cohorts, following multiallelic variant splitting and left normalization of insertion/deletions (indels) with BCFTools^60^, we removed variants with genotype quality (GQ) <20, read depth (DP) <8, call rate (CR) < 90%, Hardy-Weinberg equilibrium p<0.000001 and monomorphic sites. Then, we removed individuals with (i) >5% missing genotype call rates, (ii) cryptic relationships (pi hat >0.125) and (iii) high rates of genotype heterozygosity (>5 standard deviations). We also removed all non-European individuals as determined by principal component analysis (PCA) using SMARTPCA^61,62^ with the 1000 genome project phase 3 used as a reference population^63^. Subjects with established pathogenic variants in known Parkinson’s disease genes (*LRRK2, SNCA, PINK1, VPS35, PARK2* and *DJ-1*) were detected and excluded from in both datasets. Following QC, we obtained 1,494 IPDGC samples (1,042 PD cases, 452 controls) and 527 PPMI samples (383 PD cases, 144 controls) with a total of 366,746 and 412,223 variants respectively. We performed Transition/Transversion (Ts/Tv), Heterozygous/Homozygous (Het/Hom) and Insertion/Deletion (In/Del) ratios calculations with SnpSift V.40^64^.

### Variant annotation and selection

Variants were annotated using ANNOVAR^65^, and categorized in two ways. First, using variant frequency data within each cohort and from the gnomAD database^10^, variants were labelled according to their observed alternative allele frequency as being infrequent (<5%), rare (<1%), very rare (<0.1%), singletons (seen in a single carrier in the cohort, but also in gnomAD) and URVs (singletons absent from gnomAD). Second, using variant functional prediction categories, variants were stratified into (a) synonymous, (b) noncoding (c), nonsynonymous (NSN), (d) loss-of-function (LoF) (frameshift/non-frameshift In/Dels, stop gains/losses and splicing) and (e) protein altering (PAVs, encompassing NSN and LOF variants). We kept PAVs, NSN and LoF variants that were predicted to be damaging by the CADD algorithm^66^ (score ≥12.37). For synonymous and noncoding variants, we used variants with CADD < 12.37 as “benign” and > 12.37 as “damaging”.

### Whole-exome, gene-set and gene-wise burden analysis

All burden analyses were performed on all PD cases and controls, and then stratified for EOPD (AOO <40 years old), older than 40 years (40+ PD) and LOPD (AOO >60 years old), in order to assess the genetic profiles across different AOO groups. All burden tests were performed for each variant frequency/functional category using logistic regression with the *glm* R package^67^. Here, we modeled the variant allele counts across the entire exome for each individual with disease status, adjusting for sex, population structure (PC1-PC5) and capture metrics (10x percentage of exome coverage).

For the gene-set analysis, we used *SKAT-O* R package^11^, on the most significant whole-exome burden categories detected, and we grouped variants according to different gene-set categories. First, we used gene constraint metrics from the gnomAD database. We selected all variants within genes with high pLI, pRec and pNull scores (pLI >0.9, n=3,230; pRec >0.9, n=4,510; pNull >0.9, n=2,096). Second, we used the following gene-set definitions from the Molecular Signatures database v.6.2 (MsigDB)^12^: (a) The Hallmark gene-sets composed of 50 gene groups that summarize and represent specific well-defined biological states or processes and display coherent expression. These gene sets were generated by a computational methodology based on identifying overlaps between gene sets in other MSigDB collections and retaining genes that display coordinate expression; (b) The C2 gene-set, composed of 4,762 sets curated from various sources such as online pathway databases, the biomedical literature, and knowledge of domain experts. Third, we used gene expression data from 54 tissues from the GTEx v.8 database^14^ (gene median transcripts per million per tissue). We selected genes where expression in a given tissue was five times higher than the median expression across all tissues, as previously used^68^. We obtained SKAT-O association P-values for each gene-set after correction for multiple testing using a family-wise error rate (FWER <0.05), calculated based on 10,000 permutations.

For the gene-wise analysis, we used SKAT-O with the same covariates as the gene-set approach and correcting for multiple testing using FWER < 0.05. Genes with at least two carriers and an uncorrected P-value of <0.05 were kept for further PPI network and single-cell RNASeq enrichment analyses.

### Protein-protein interaction network analysis and single-cell enrichment of gene-wise burden candidates

To study the interplay between proteins coded by genes prioritized by the gene-wise burden analysis, we utilized the R package WebgestaltR^15^ to build PPI networks via network topology analysis and random walk algorithm. Using the Network Retrieval and Prioritization mode, we submitted the list of genes in different combinations: first, the gene-wise burden candidates list alone (**Supplementary Table 13**), then adding 26 known and suggested EOPD genes (**Supplementary Table 15**) and finally including 303 genes of interest tagged by the 90 risk loci coming from the recent PD GWAS meta-analysis^1^. GO analysis enrichment were obtained from the Biogrid PPI Networks database^69^. FDR adjustment was used to correct for multiple testing. In order to obtain PPI enrichment statistics, we submitted the same combination of genes to STRING^16^, and retrieved high confidence networks. For visualization and feature annotation purposes we loaded the resulting networks to Cytoscape v.3.7.2^70^.

To observe if genes coming from the gene-wise burden analyses play a role in specific brain cell types, we used the Expression Weighted Cell Type Enrichment analysis (EWCE)^18^ with single-cell and single-nuclei RNASeq expression datasets from 24 brain cell types from mice^55^. Briefly, the EWCE pipeline identifies mouse orthologs (if available), and then tests whether genes have higher expression levels in a given cell type than can reasonably be expected by chance. The enrichment tests are corrected using the Benjamini-Hochberg FDR method.

### Heritability estimation of ultra-rare variants

We used the Singleton Heritability inference with REML (SingHer) R package^20^ to calculate the heritability of URVs in EOPD, all PD cases and 40+ in the IPDGC cohort.

## ACKNOWLEDGEMENTS

We would like to thank the all the patients that provided their genetic information. For IPGCG acknowledgements please visit https://pdgenetics.org/partners.

## AUTHOR CONTRIBUTIONS

B.I.B. and S.J.L. conceived and designed the experiments. B.I.B and S.J.L. performed experiments or analyzed the data. D.K. and IPDGC members helped in drafting the manuscript. All authors reviewed the manuscript for scientific accuracy.

## DATA AVAILABILITY

IPDGC and PPMI whole-exome sequencing data are publicly available upon application through their respective project websites (https://pdgenetics.org/resources, http://www.ppmi-info.org/access-data-specimens/download-data/)

