## Supplementary Information for "Whole-exome analysis in Parkinson’s disease reveals a high burden of ultra rare variants in early onset cases"

### Supplementary Figures

**Supplementary Figure 1.** Study pipeline.

**Supplementary Figure 2.** Whole-exome burden of synonymous and noncoding variants in IPDGC.

**Supplementary Figure 3.** Whole-exome burden in the PPMI cohort.

**Supplementary Figure 4.** Whole-exome burden of synonymous and noncoding variants in the PPMI cohort.

**Supplementary Figure 5.** Gene-set burden analysis in the IPDGC PD cases over 40 years old.

**Supplementary Figure 6.** Protein-protein Interaction network for gene-wise burden of novel-singleton variants in EOPD.

**Supplementary Figure 7.** Protein-protein Interaction network for gene-wise burden candidates and GWAS PD risk genes.

### Supplementary Tables (Excel)

**Supplementary Table 1.** Average alternative-allele counts for exome-wide variants according to frequency and functional classification in IPDGC and PPMI datasets.

**Supplementary Table 2.** Burden results for IPDGC dataset.

**Supplementary Table 3.** Burden results removing lower frequency variants for IPDGC dataset.

**Supplementary Table 4.** Burden results for synonymous and noncoding variants in the IPDGC dataset.

**Supplementary Table 5.** Burden results URVs with depth > 20 in the IPDGC dataset.

**Supplementary Table 6.** Burden results for PPMI dataset.

**Supplementary Table 7.** Burden results without URVs for PPMI dataset.

**Supplementary Table 8.** Burden results of synonymous and noncoding variants for the PPMI dataset.

**Supplementary Table 9.** Burden analysis variants on gene constraint gene-sets for 40+ PD and LOPD in the PPMI dataset.

**Supplementary Table 10.** MsigDB gene-set SKAT-O results for the PPMI variants.

**Supplementary Table 11.** SKAT-O burden test for the lysosomal storage disorder and mitochondrial DNA maintenance pathways previously associated with PD.

**Supplementary Table 12.** GTEx v8 gene-set SKAT-O results for the PPMI variants.

**Supplementary Table 13.** Gene-wise SKAT-O nominal P-value results for IPDGC EOPD PAV variants.

**Supplementary Table 14.** Gene-ontology enrichment for PPI network from 308 URV harboring genes.

**Supplementary Table 15.** List of known and suggested EOPD genes and their model of inheritance.

**Supplementary Table 16.** Gene-ontology enrichment for PPI network from 308 URV harboring genes and 26 known and suggested PD genes.

**Supplementary Table 17.** Gene-ontology enrichment for PPI network from 308 URV harboring genes and 352 candidate genes tagged by the 90 PD risk loci from GWAS meta-analysis.

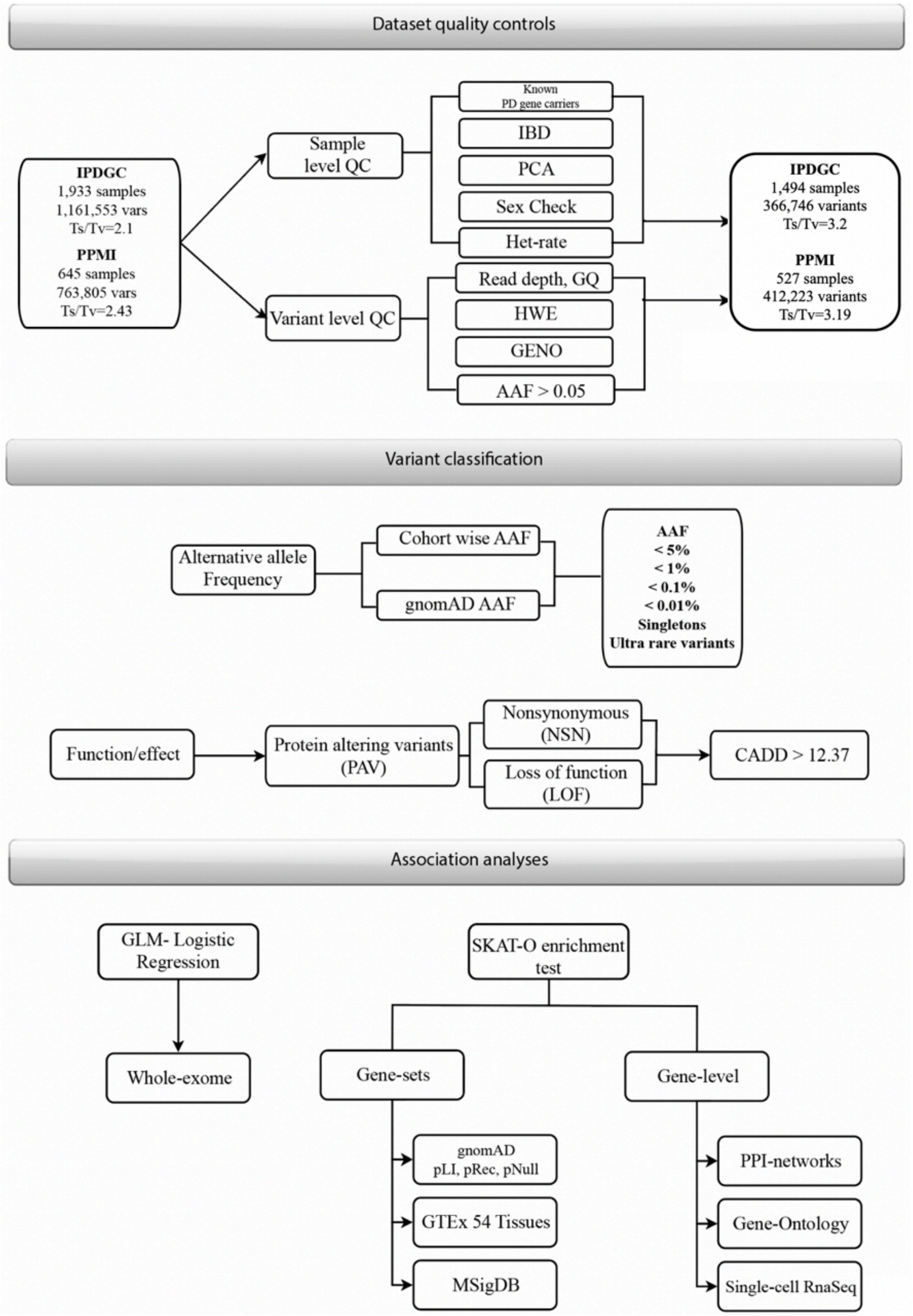


**Supplementary Figure 1. Study pipeline**. Main procedures performed in the study are depicted in the flowchart as follows: Dataset quality controls, Variant classification and association analyses. IPDGC: International Parkinson's Disease Genomics Consortium. PPMI: Parkinson's Progression Marker Initiative. TsTv: Transition Transversion Ratio. IBD: Identity by descent. PCA: Principal Component Analysis. GQ: Genotype Quality. HWE: Hardy-Weinberg Equilibrium. GENO: Genotype missingness. AAF: Alternative Allele Frequency. CADD: Combined Annotation Dependent Depletion. GLM: Generalized linear models. SKAT-O: Optimized Sequence Kernel Association Test. pLI, pRec, pNull: gnomAD gene constraint metrics. GTEx: Genotype-Tissue Expression project. MSigDB: Molecular Signature database. PPI: Protein-Protein Interactions.

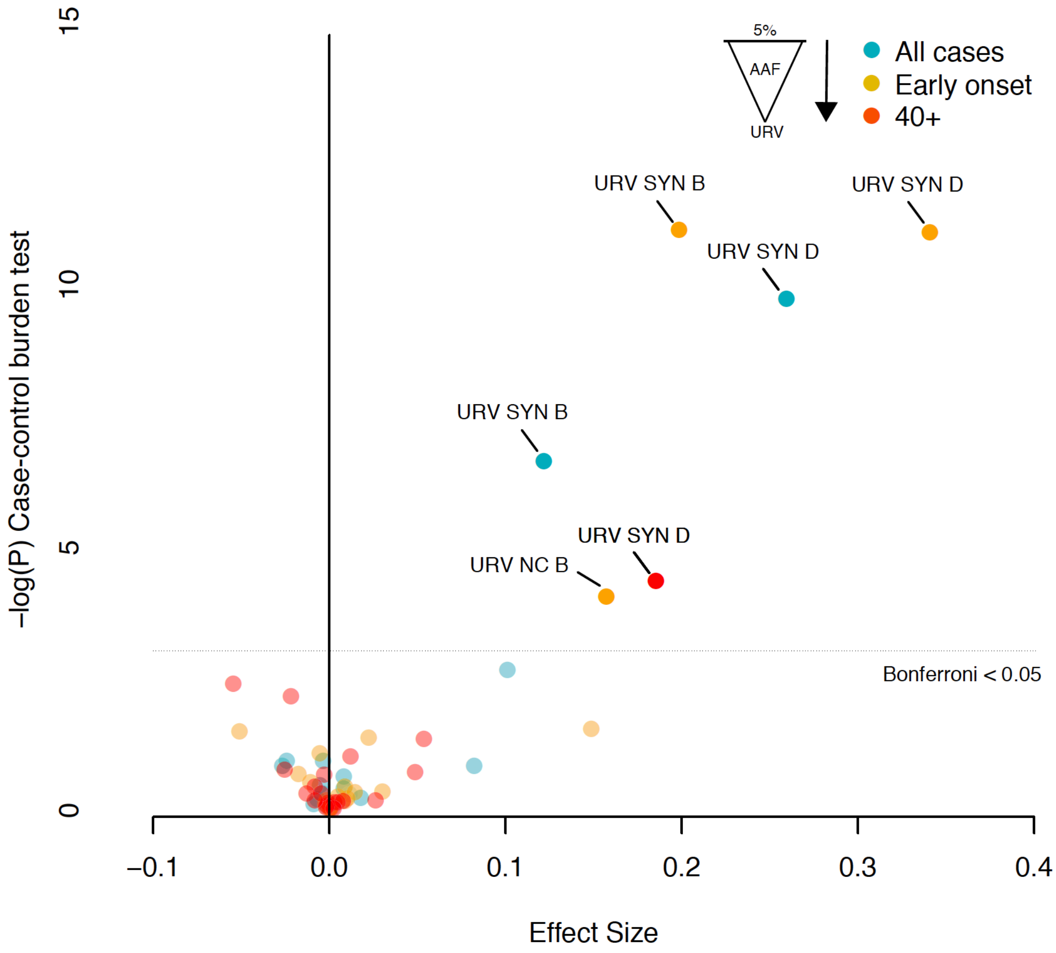


**Supplementary Figure 2. Whole-exome burden of synonymous and noncoding variants in IPDGC**. Burden analysis of different variant frequencies and functional categories for all PD cases, cases over 40 (40+) years old and over 60 years old (LOPD). SYN: synonymous. NC: non-coding. B: benign variants with CADD < 12.37. D: Damaging variants with CADD > 12.37. AAF: Alternative allele frequency; PAV: Protein-altering variants; LOF: Loss of function variants.

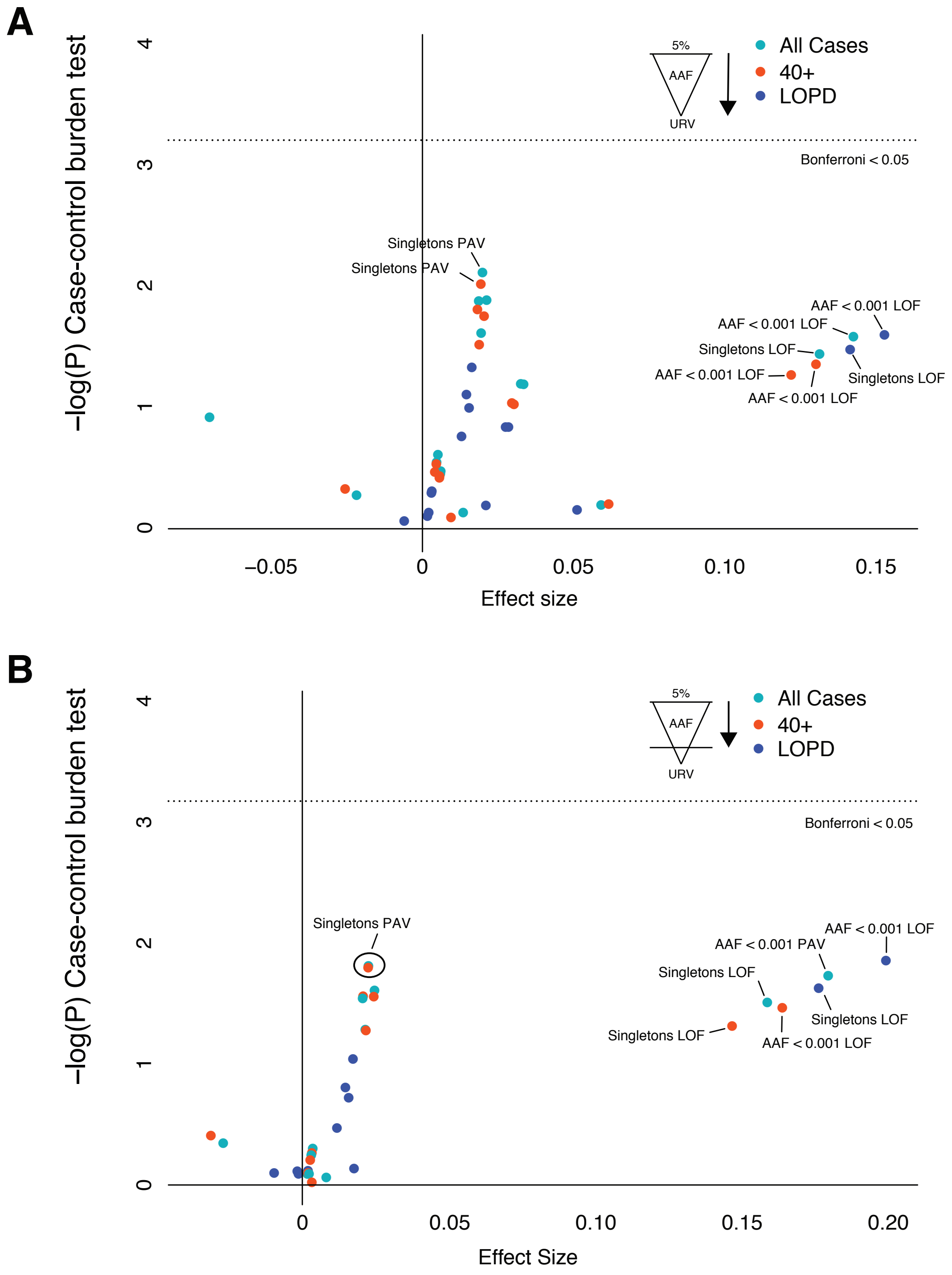


**Supplementary Figure 3. Whole-exome burden in the PPMI cohort**. Burden analysis of different variant frequencies and functional categories for all PD cases, cases over 40 (40+) years old and over 60 years old (LOPD). **A)** Burden towards rarer variants from 5% to URVs. **B)** Burden towards rarer variants removing URVs from each variant category. AAF: Alternative allele frequency; PAV: Protein-altering variants; LOF: Loss of function variants

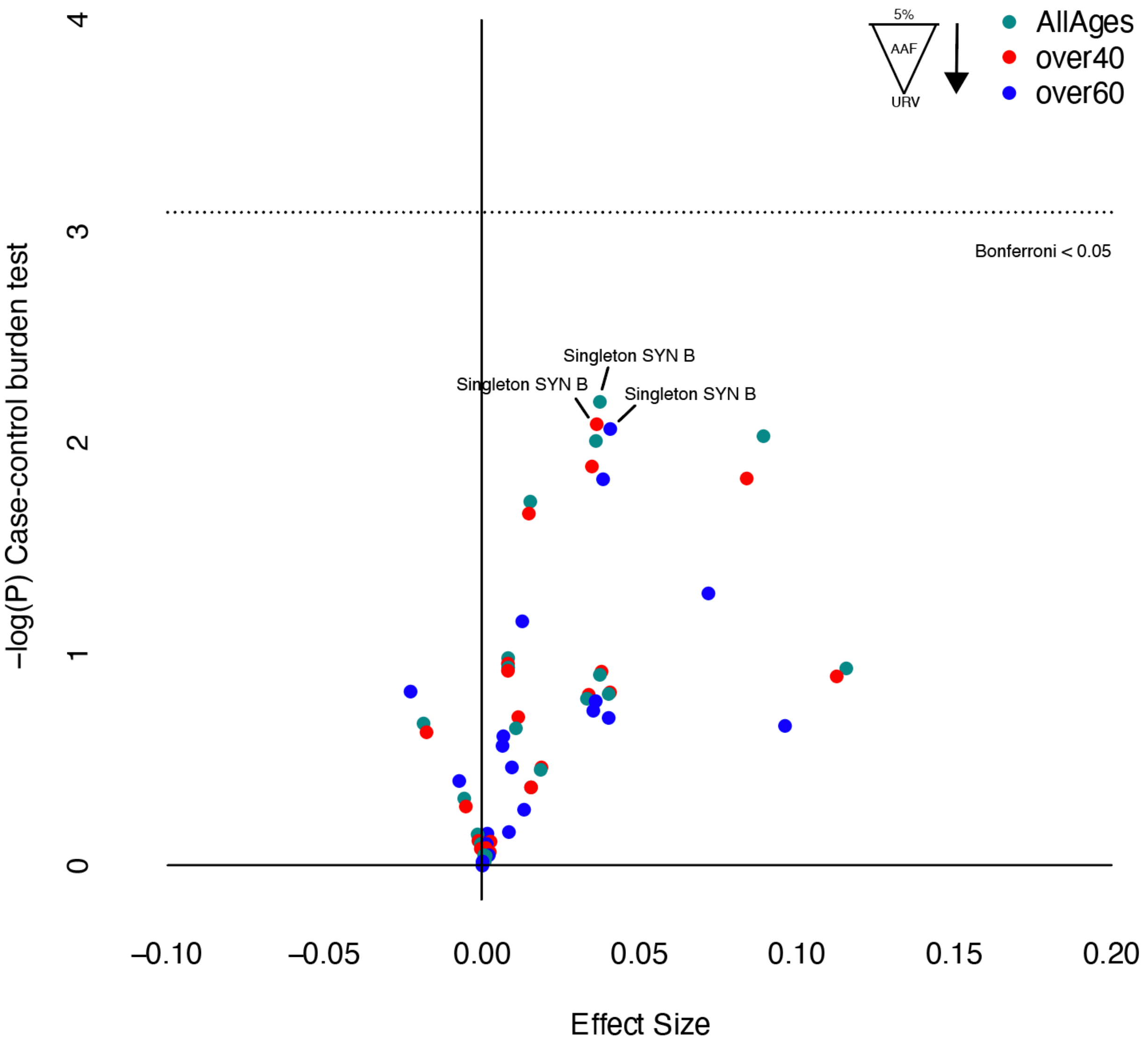


**Supplementary Figure 4. Whole-exome burden of synonymous and noncoding variants in the PPMI cohort**. Burden analysis of different variant frequencies and functional categories for all PD cases, cases over 40 (40+) years old and over 60 years old (LOPD). SYN: Synonymous variant; B: benign variants with CADD < 12.37.

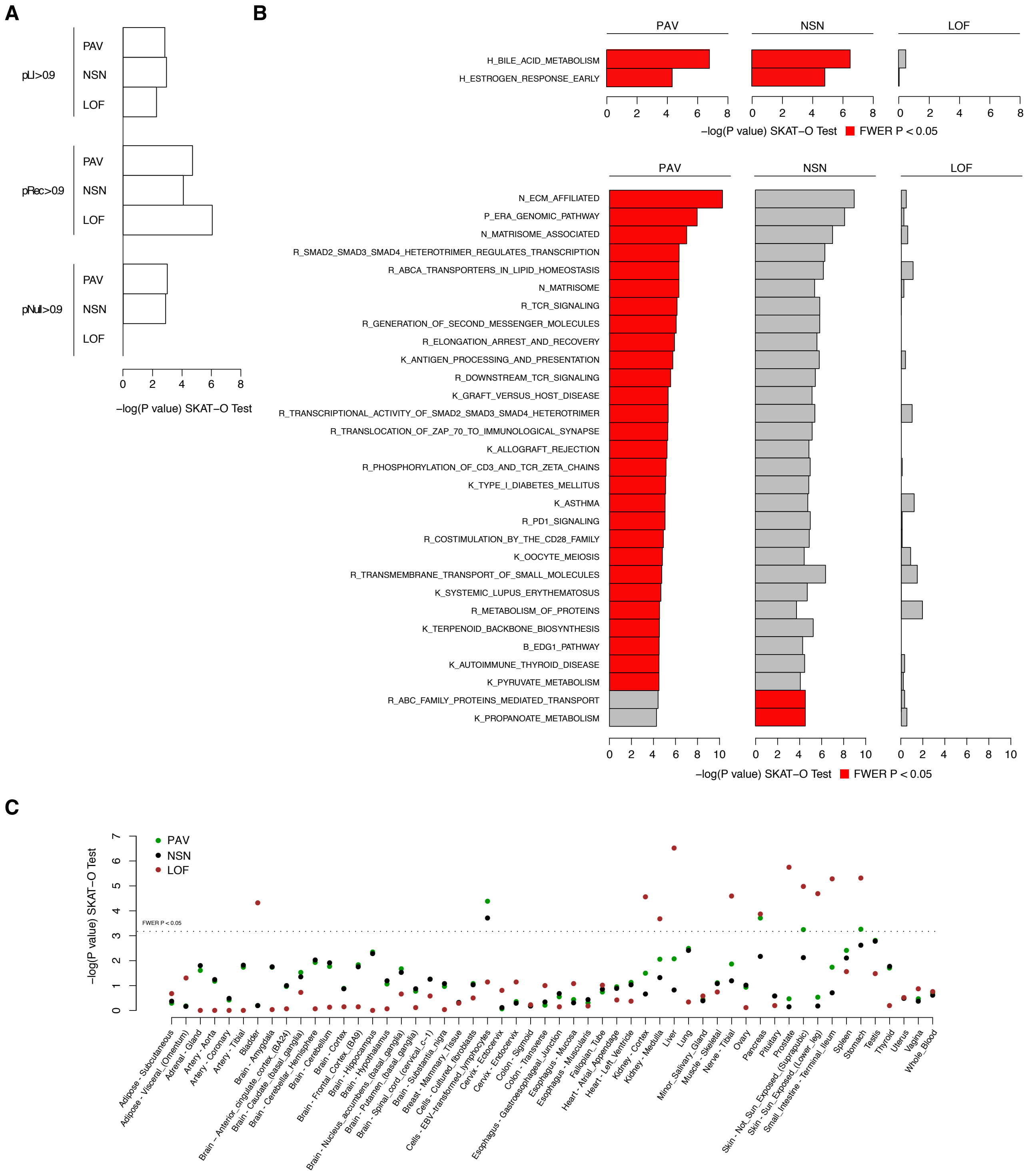


**Supplementary Figure 5. Gene-set burden analysis in the IPDGC PD cases over 40 years old**. Gene-set analysis performed with SKAT-O using PAVs, NSNs and LOFs variants with AAF > Singleton using gene lists from: **A)** gnomAD gene constraint metrics pLI, pRec and pNull > 0.9; **B)** MsigDB Hallmark pathways (upper panel) and C2 curated pathways (lower panel); **C)** High expression genes from 54 tissues in the GTEx database V8. Family-wise error rate P < 0.05 was used as a threshold for statistically significant results.

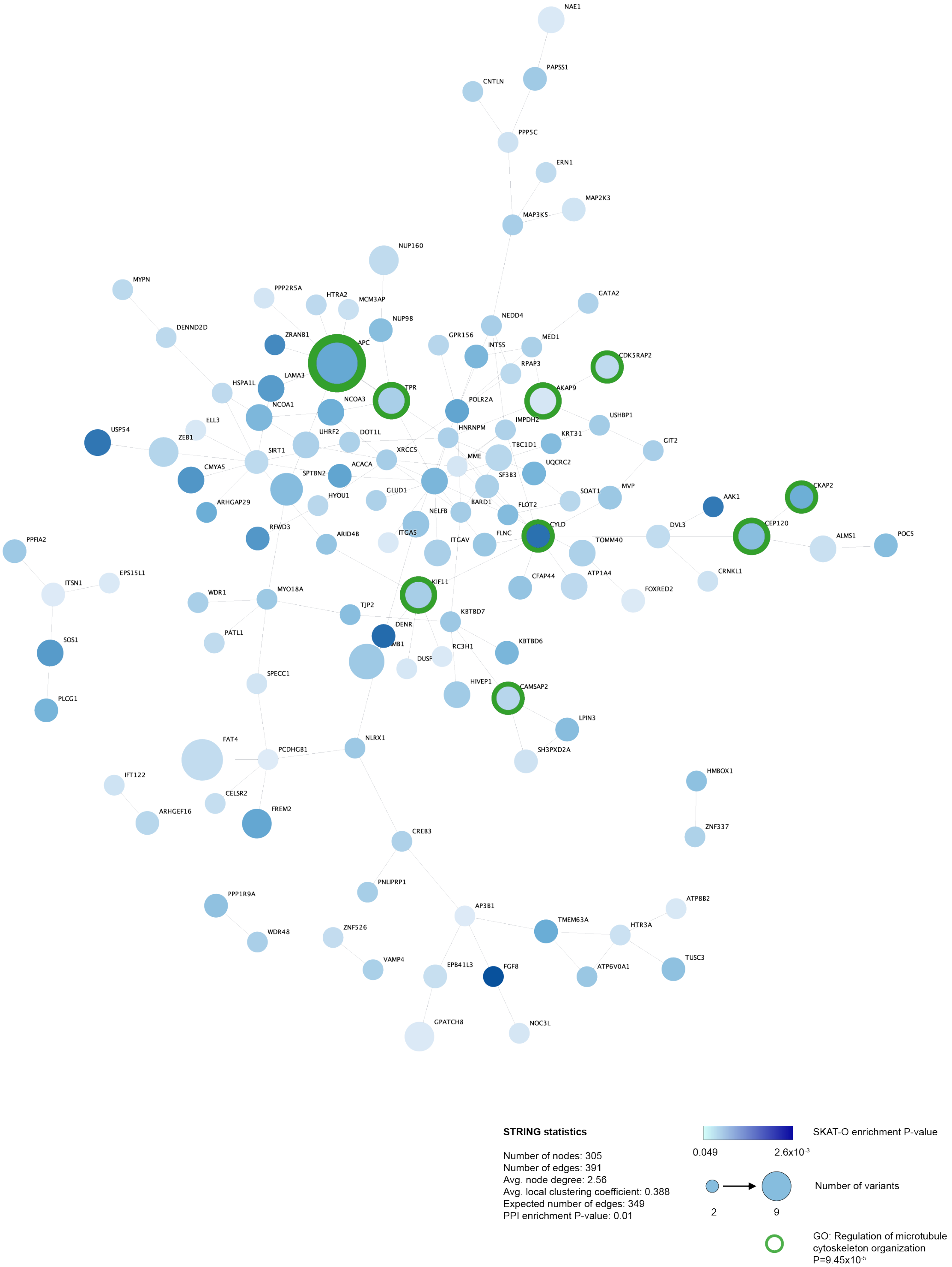


**Supplementary Figure 6. Protein-protein Interaction network for gene-wise burden of novel-singleton variants in EOPD**. Significant network built using 308 genes with SKAT-O P < 0.05 from EOPD URVs with WebGestalt Network Topology-based Analysis (NTA). Statistics from STRING using the same list of genes are shown. Shades of blue indicate SKATO P-values for less (light blue) to more significance (dark blue). Node size proportionally represent the number of variants observed in each gene. The green perimeter shows genes member significant gene-ontology enrichment results.

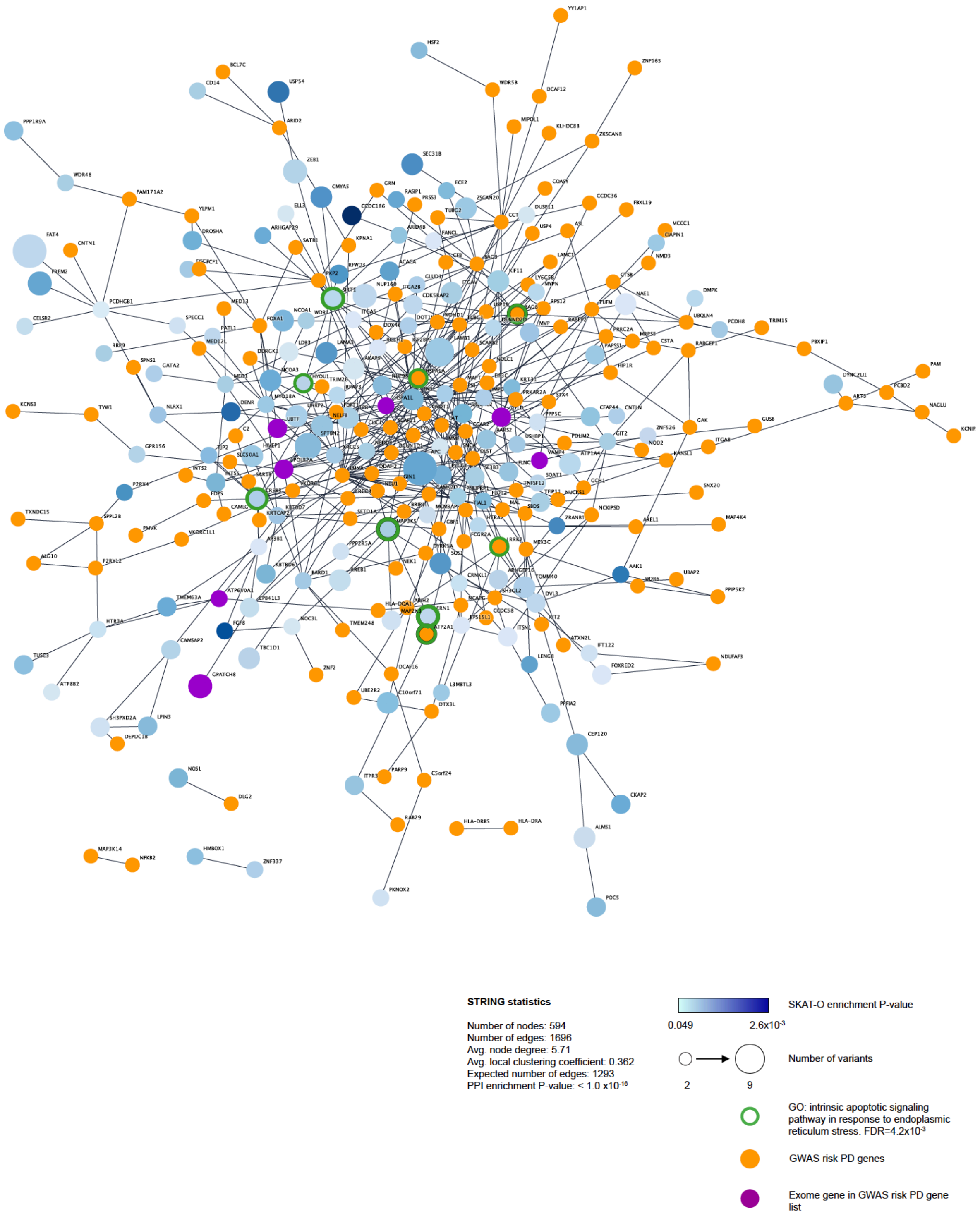


**Supplementary Figure 7. Protein-protein Interaction network for gene-wise burden candidates and GWAS PD risk genes**. Significant network using the list of 308 genes coming from the SKAT-O test with $P < 0.05$ (EOPD URVs) plus 303 genes from a recent PD GWAS meta-analysis (611 genes total, 594 mapped to STRING DB). Shades of blue indicate SKATO P-values for less (light blue) to more significance (dark blue). Node size proportionally represent the number of variants observed in each gene. The green perimeter shows genes member significant gene-ontology enrichment results. Nodes in orange represent genes coming from the GWAS study, purple nodes represent overlaps of gene-wise burden candidates with genes present in the GWAS results.
